# Coupling and Heterogeneity Modulate Pacemaking Capability in Healthy and Diseased Two-Dimensional Sinoatrial Node Tissue Models

**DOI:** 10.1101/2022.04.14.488275

**Authors:** Chiara Campana, Eugenio Ricci, Chiara Bartolucci, Stefano Severi, Eric A. Sobie

## Abstract

Both experimental and modeling studies have attempted to determine mechanisms by which a small anatomical region, such as the sinoatrial node (SAN), can robustly drive electrical conduction in the human heart. However, despite important advances from prior research, important questions remain unanswered. This study aimed to investigate, through mathematical modeling, the role of intercellular coupling and cellular heterogeneity in synchronization and pacemaking within the healthy and diseased SAN. In a multicellular computational model of a monolayer of either human or rabbit SAN cells, simulations revealed that heterogenous cells synchronize their discharge frequency into a unique beating rhythm across a wide range of cellular heterogeneity and intercellular coupling. However, an insightful and unanticipated behavior appears in pathological conditions precipitated by perturbations of certain ionic currents (g_CaL_ = 0.5): an intermediate range of intercellular coupling (900-4000 MΩ) is beneficial to the human SAN automaticity, enabling a very small portion of tissue (3.4%) to drive propagation, which fails with both lower and higher resistances. This protective effect of intercellular coupling and heterogeneity, seen in both human and rabbit tissues, highlights the remarkable resilience of the SAN. Overall, the model here developed allowed insight into the mechanisms of automaticity of the human sinoatrial node. The simulations suggest that certain degrees of gap junctional expressions protect the SAN from ionic perturbations that might be caused by drugs or mutations.

## INTRODUCTION

Understanding the mechanisms that coordinate the spontaneous firing of the sinoatrial (SA) node has long been an issue of great interest in cardiac electrophysiology. After early studies believed that a single pacemaker region drives the entire SA node, more recent research has shown that the heartbeat originates from the coordination of a complex structure [1]. Many studies have worked to unravel the basis of this coordination, through both experiments [2–4] and mathematical modeling [5–7]. Despite the many insights obtained by these studies, important questions remain unresolved, particularly with respect to how heterogeneity between SA nodal myocytes and inter-cellular coupling combine to influence coordinated beating in tissue. For example, althought it has recently been shown experimentally that not all SA nodal cells fire spontaneously when they are enzymatically isolated [8], we do not know how non-firing cells when they are electrically coupled in tissue, nor how the percentage of non-firing cells influences the overall behavior of the SA node.

Multiple mathematical models exist in the literature that describe the behavior of isolated SA nodal myocytes. Most of these have been developed on the basis of data obtained in animal models, especially rabbits [9,10], but a model based on human data has been published more recently [11]. Although it is obviously helpful to have multiple tools available for computational analyses, a question that commonly arises in such circumstances is the extent to which the behavior observed in a particular model is generalizable. On the other hand, when similar trends are seen across multiple mathematical representations, this can provide confidence in the model predictions.

In this study, we studied through cellular and tissue simulations how heterogeneity between SA nodal mycoytes and intercellular coupling influence the coordination of beating within the SA node. The main goals were to: i) assess the effect of cellular heterogeneity in isolated SA nodal cells; ii) gain mechanistic insight into how electrical coupling between SA nodal cells modulates pacemaker activity at different levels of heterogeneity; iii) investigate the consequences of simulated diseased condition on SAN autorhythmicity. Heterogeneous populations of SA nodal myocytes were generated at several levels of variability, and physiological behavior was simulated in both isolated cells and 2-dimensional tissue. Major results of the simulations were: i) cellular heterogeneity increases AP frequency and duration as well as the percentage of “dormant” cells, with remarkable consistency between three SA nodal myocyte models [9–11]; ii) intercellular coupling allows the cells to synchronize the beating rate in all conditions, except when heterogeneity is large and coupling between myocytes is waek; iii) blockade of particular ionic currents leads to a loss of robustness in which coordinated beating of the tissue fails at high and low coupling but can be maintained within a narrow range of intermediate coupling values. Overall, these simulations provide insight into the conditions that promote synchronized beating in the SA node, and how this can be maintained in the presence of heterogeneity.

## METHODS

### Study Design

The goal of our study was to analyze, through mechanistic simulations, how heterogeneity between cells and gap junctional coupling influence the automaticity of the sinoatrial node and entrainment of the Action Potential (AP). As schematically shown in Figure 1, a two-dimensional tissue model was developed starting from models of the isolated sinoatrial node (SAN) cell of different species (human and rabbit). For human SAN, the recent Fabbri et al. [11] model was used, whereas for rabbit SAN, both Maltsev-Lakatta [9] and Severi [10] models were considered (Figure 2). Such models can be utilized to determine the effect of structural remodeling due to conditions such as low coupling (mimicking diffuse fibrosis) and clusters of non-spontaneous (“dormant”) cells.

**Figure 1.**
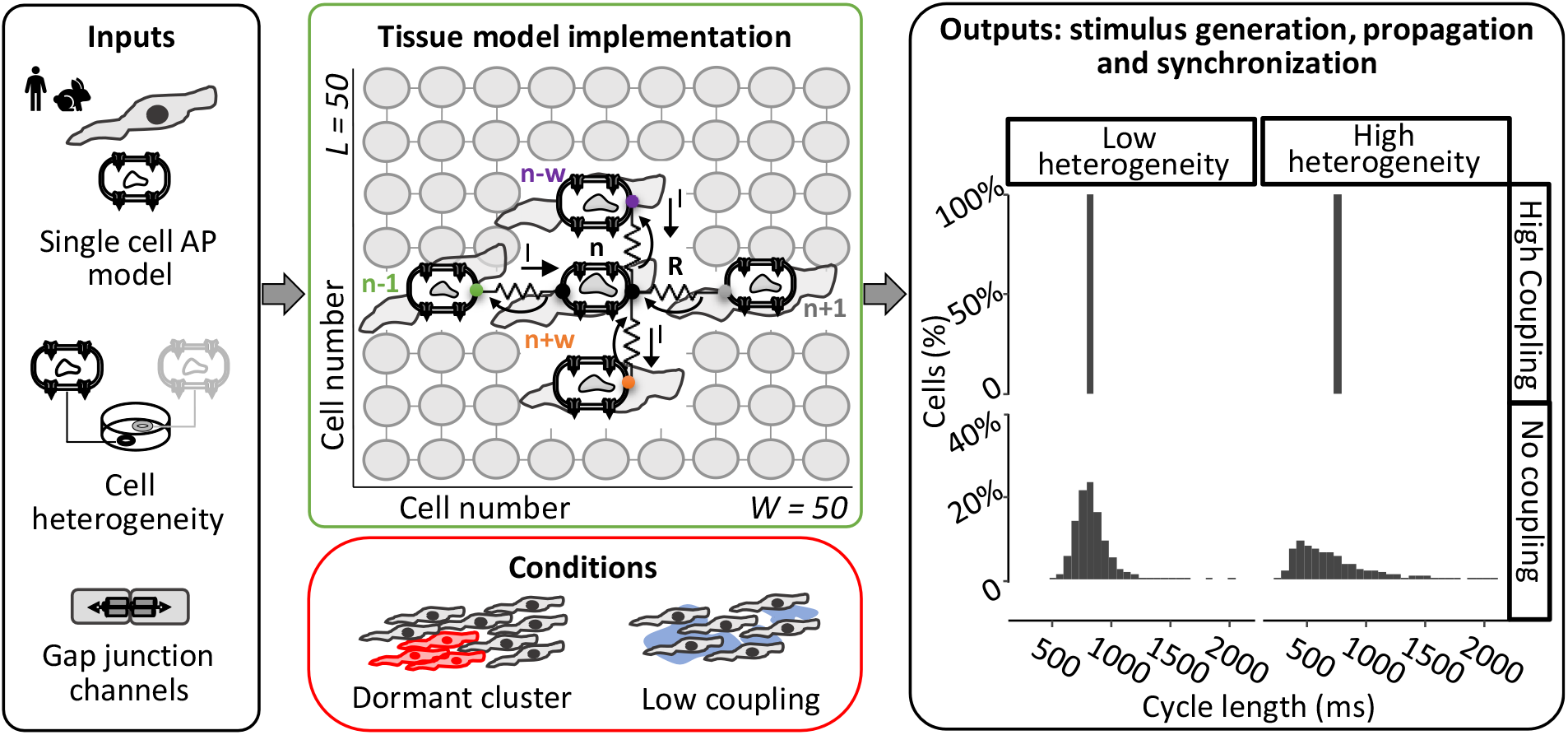
Schematic of multicellular study design. A multiscale mathematical modeling approach was formulated and adopted to study the mechanisms of sinoatrial node excitability. The effect of cell-to-cell coupling and cellular heterogeneity on tissue synchronization were evaluated both in the healthy and diseased sinoatrial node.

**Figure 2.**
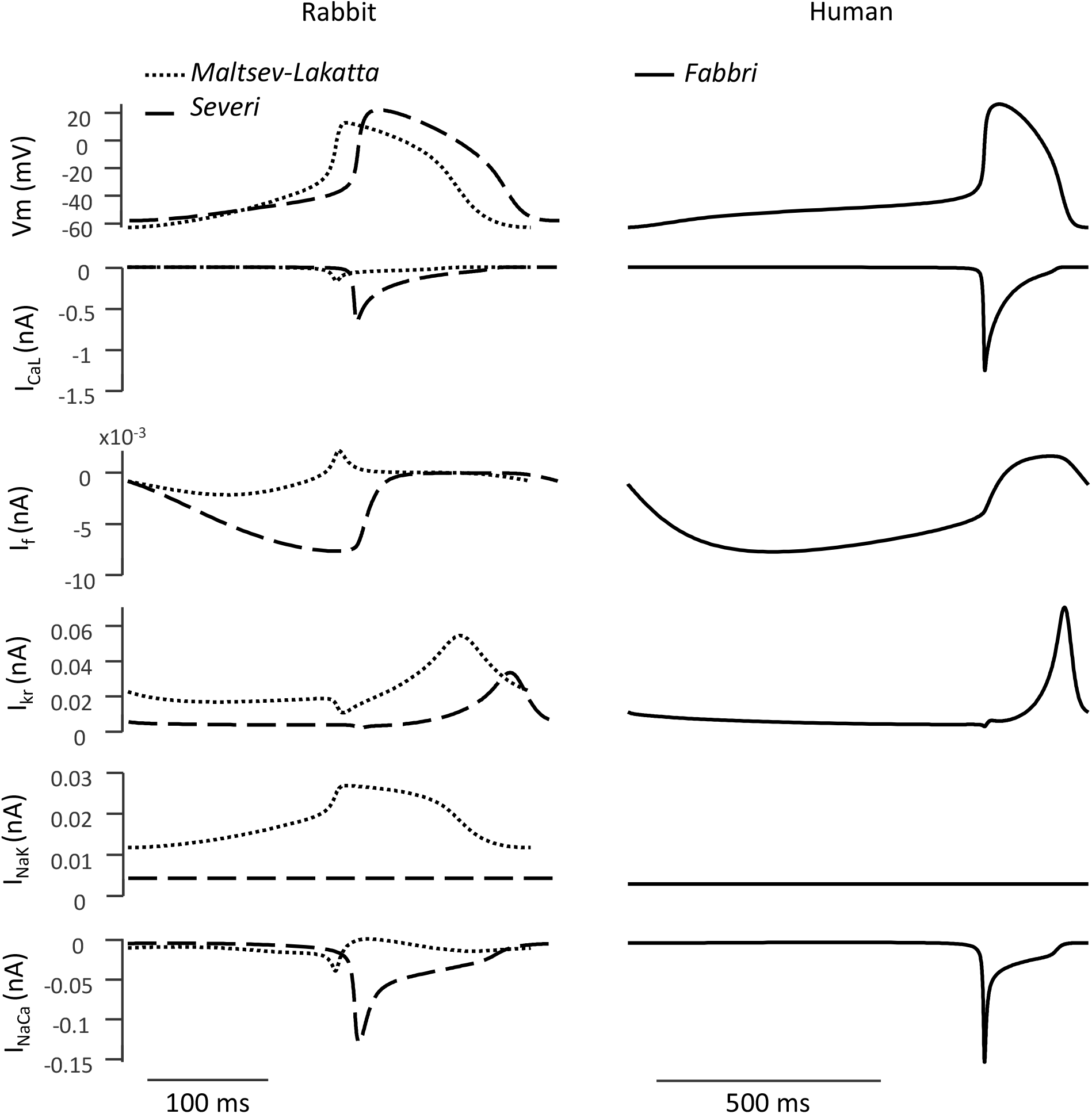
Simulation of rabbit and human sinoatrial node action potential and ionic currents. Three models were used to simulate the spontaneous pacemaker activity of the sinoatrial node [9–11]. The figure shows how the AP traces and selected ionic currents differ among the three models. The main differences are: (1) the diastolic depolarization (DD) phase, hence the cycle length is much longer in human than in rabbit models, and (2) as expected, the funny current has a prominent role in the DD phase of Severi and Fabbri models, while it is less important in the Maltsev-Lakatta model.

### Modeling heterogeneous populations of SAN cells

Cellular heterogeneity was simulated with each model by varying the maximal conductance of the ionic channels. The baseline conductance value of each channel was multiplied by a random scale factor chosen from a a lognormal distribution [12,13]. Five different values of the shape factor (σ) of the lognormal distribution (σ from 0.1 to 0.5) were used to account for different levels of heterogeneity. The purpose of creating heterogeneous populations of cells was three-fold. First, we used these populations to run a sensitivity analysis where the contributions of individual ionic currents on the cell’s automaticity were evaluated with a logistic regression model [14]. Second, isolated cell simulations were run to assess the effects of σ on the AP parameters and on each model’s robustness (that is, how many cells showed spontaneous beating after parameter randomization). Third, the cellular populations were used to create the two-dimensional propagation model in an attempt to recapitulate part of the complexity and heterogeneity characteristic of the SAN structure. In particular, we aim to compare the behavior of isolated and coupled cells to gain a mechanistic understanding of how coupling modulates the effects of heterogeneity.

### Mathematical modeling of electrical propagation throughout the SAN

We implemented a tissue model by connecting individual SAN cells through an intercellular resistance that represents the gap junctional channels. In this model, each cell is described by a system of ordinary differential equations that, integrated over time, yields the value of ionic concentrations and gating variables (state vector). In addition, the membrane potential is calculated through a partial differential equation since its value depends both on the individual cell and the neighboring cells in the tissue. Thus, the updating of the membrane potential is described by the following equation:

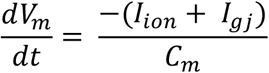

where *V*_*m*_ is the membrane potential, *C*_*m*_ is the cellular capacitance, *I*_*ion*_ is the sum of all the ionic currents (dependent on the model) and *I*_*gj*_ is the sum of the currents exchanged with the four neighboring cells. We define the sign of I_gj_ such that negative I_gj_ represents current flowing into a particular cell from its neighbors, which depolarizes that cell. Importantly, this model was implemented such that the state vector is computed in parallel for all cells in the tissue, which substantially decreases the execution time and allowed us to run numerous simulations to test different hypotheses. Full details on the hardware and software specifications are provided in the Supplementary Material.

### Simulation protocols and conventions for model outputs

We considered a tissue formed of 2500 cells of equal size (capacitance), arranged in a 50 × 50 matrix. Simulations were executed for a duration of 20 s. In addition to different amounts of cellular heterogeneity, we tested multiple levels of intercellular coupling from a resistance value of 10 MΩ (100 nS; strongly coupled cells) to 10,000 MΩ (0.1 nS; weakly coupled cells) [15,16]. The outputs of these simulations for each cell in the tissue were: membrane potential (V_m_), ionic current of each SAN cell (I_ion_), and gap junctional current (I_gj_). Additionally, I_net_ is defined as the sum of I_gj_ and I_ion_, reflecting the total net current of each cell. A negative I_net_ depolarizes the membrane, whereas a positive I_net_ hyperpolarizes it.

From the AP trace (Figure 3) we defined MDP (mV) as the maximum absolute value of voltage during diastolic depolarization; OS (mV), or overshoot, as the maximum value of voltage; take-off potential, or TOP (mV), as the voltage at the first time step during diastolic depolarization when 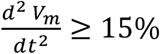 of the maximum 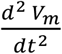 (Kohajda et al., 2020). These three outputs were then used to compute the metrics on which our analysis relied: DD (ms), or diastolic depolarization, is the phase of the AP between MDP and TOP; APD (ms), or action potential duration, is the time difference between TOP and the following MDP; CL (ms), or cycle length, is the time difference between two consecutive OS; APA (mV), or action potential amplitude, is the difference in voltage between OS and MDP. Cells were considered as having the property of automaticity when they generated at least 3 peaks (OS ≥ 0 mV; MDP ≤ -40 mV) in the final 5 s of simulation.

**Figure 3.**
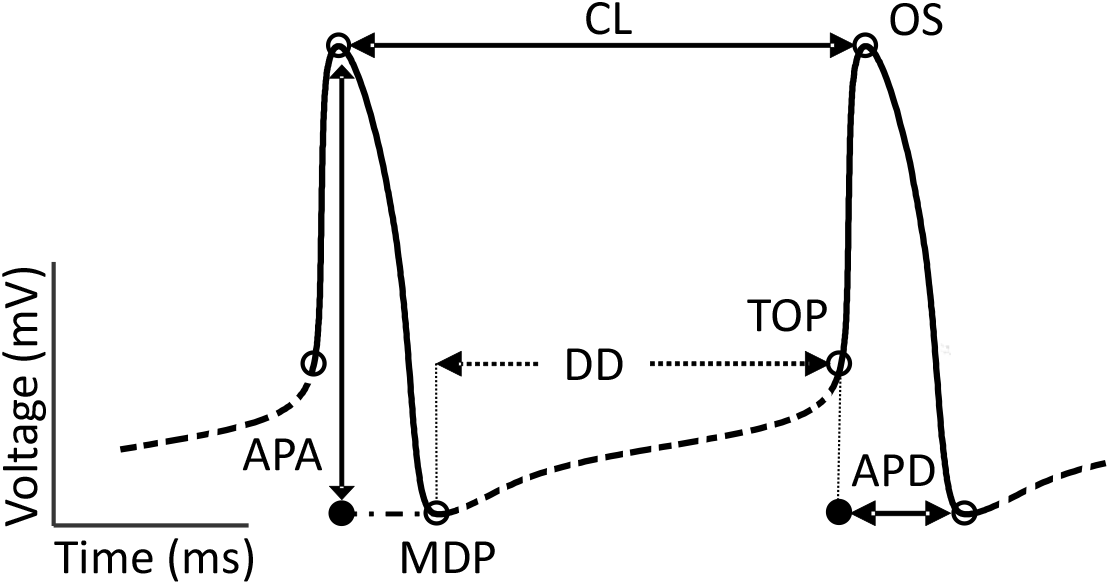
Features extracted from sinoatrial node AP simulations. A schematic AP trace is annotated with characteristic features. Details of the calculations are outlined in the text. Dashed line indicates the diastolic phase (DD). Solid line indicates AP phase.

### Categorization of cells inside the tissue

To better describe their behavior, the cells forming the 2D tissue were divided into categories. Initially, cells were defined as “spontaneous” or “dormant” depending on whether they showed rhythmic electrical activity when simulated in an uncoupled condition (R = ∞ MΩ). When a mixture of spontaneous and dormant cells is coupled in tissue, the dormant cells may exhibit action potentials under many conditions. Understanding this concept requires the definition of subcategories, as illustrated in Figure 4, that capture different types of cellualr behaviour.

**Figure 4.**
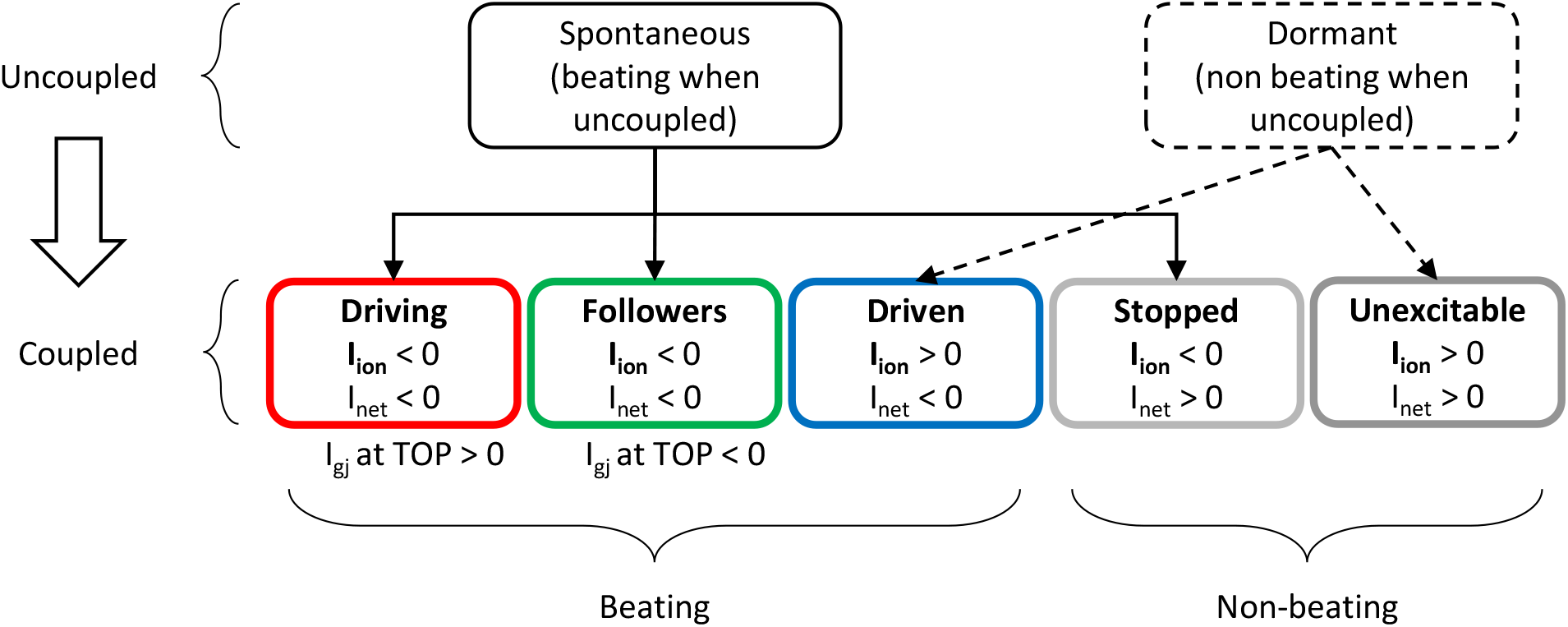
Cell categorization. Cells forming the tissue have been divided into different categories wheter they were beating or not in both coupled and uncoupled conditions.

As schematized in Figure 4, based on their behavior when coupled within the tissue, “spontaneous” isolated cells could be further classified into: (1) “driving” if they continued to show rhythmic APs and had a positive (outward) I_gj_ at TOP. This meant that they reached threshold before adjacent cells and delivered current to their neighbors; (2) “followers” if, in spite of their spontaneous activity when uncoupled, they had a negative inward I_gj_ at TOP in the coupled condition, meaning that adjacent cells supplied current to assist their depolarization; (3) “stopped” if they did not show APs. On the other hand, “dormant” cells showed two different behaviours when coupled: (1) isolated “dormant” cells that started to beat thanks to coupling were called “driven,” whereas (2) cells that remained silent were termed “unexcitable.” Note that since “unexcitable” cells do not show APs under any condition, features such as TOP and DD are undefined for these cells. For “stopped” cells we calculated DD and TOP based on simulations performed in the uncoupled condition. This procedure allowed us to investigate the current generated and exchanged at corresponding time points when they were coupled.

## RESULTS

### Increased heterogeneity causes failure of spontaneous beating in a fraction of isolated SAN cells

The behavior of the single SAN cell models used in this work is shown in Figure 2. Following the approach described in the Methods, we introduced heterogeneity in the ionic currents underlying the AP of each model. Figure 5 illustrates the impact of heterogeneity on the excitability and electrical properties of the isolated SAN cells. It is evident from Figure 5A that at increasing levels of the heterogeneity factor σ, several cells lose their automaticity. The exact percentage of cells that fails to depolarize spontaneously depends on the model. The Severi model is very resistant to parameter changes, while Fabbri and Maltsev-Lakatta models are more susceptible to this heterogeneity. In Figure 5B the AP metrics are summarized for the cells that retain their automaticity throughout various levels of heterogeneity. Across all models, at increasing heterogeneity, variability in amplitude, duration and frequency of the AP increases. Additionally, the Fabbri model shows a decrease in the mean value of cycle length at increasing heterogeneity.

**Figure 5.**
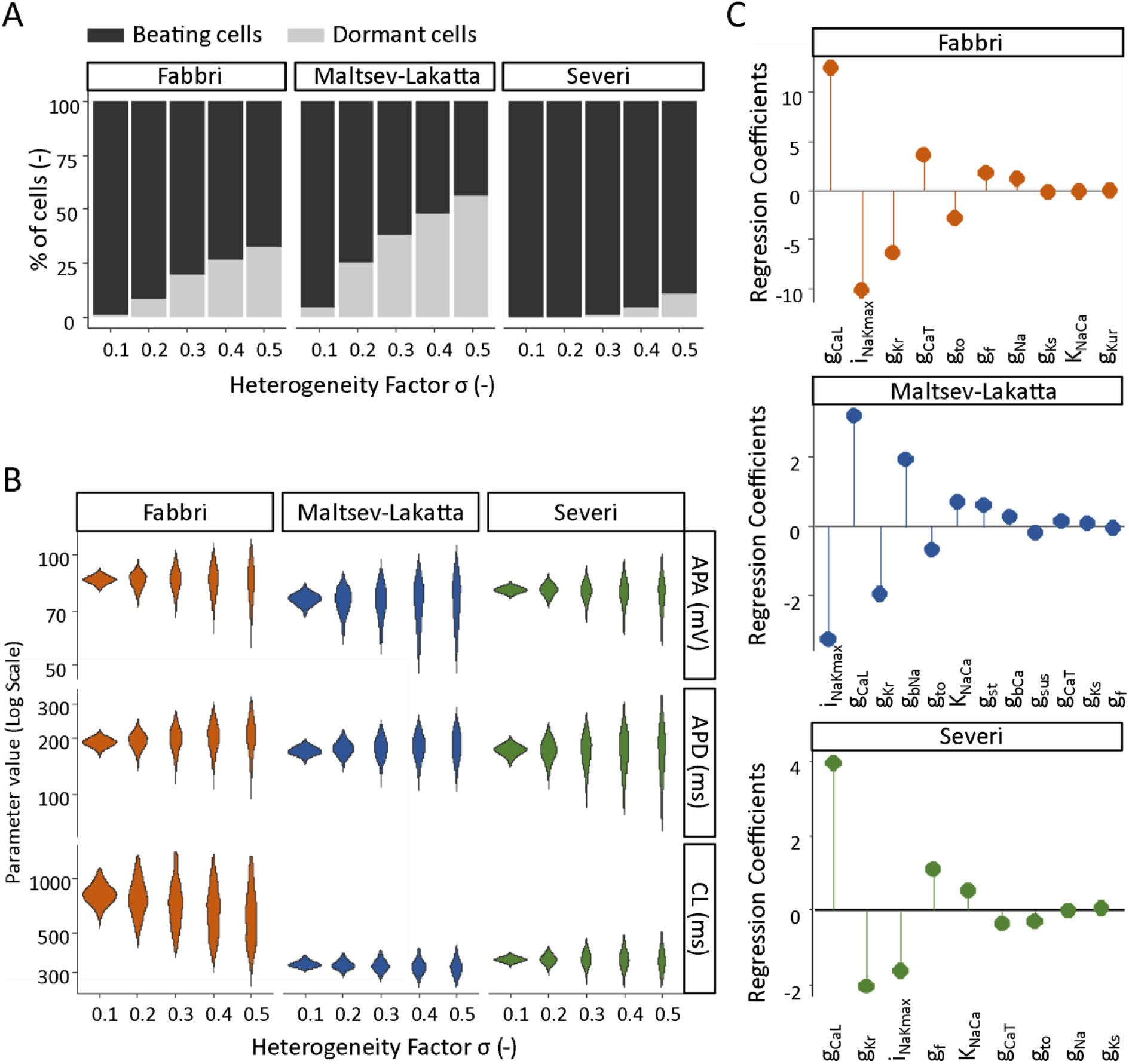
Modeling conductance heterogeneity using virtual populations of single SAN cells. (A) The effect of heterogeneous ionic channel expression on the automaticity of SAN cells was compared across models. In all three models the percentage of cells that became silent rose at increasing amounts of heterogeneity. (B) The effect of heterogeneity on the SA node AP properties was evaluated in spontaneously beating cells, by measuring cycle length (CL), AP amplitude (APA) and AP duration (APD) at varying σ levels. (C) Logistic regression analysis was utilized to deduce which specific ionic currents across the three models are responsible for SA node cell’s automaticity. Positive values indicate that an increase in the parameter increases the probability of the cell to be spontaneously beating.

We interpreted this result to mean that when we increase the heterogeneity, a certain number of cells beat at a decreased frequency until they stop all together and the residual cells remaining active were the faster beating cells. In Figure 5C, we examined which specific ionic currents were responsible for the automaticity. We calculated the regression coefficients of the logistic regression model and found that I_CaL_ was the most important variable and correlated with an increased probability of automaticity across all three models. Conversely, I_Kr_ and I_NaK_ were correlated with the decreased probability of spontaneous beating. g_bNa_ stands out as a difference between models: it is the second most important inward current in Maltsev-Lakatta model (despite it being a background current), but is completely absent from the other two models.

Finally, it should be highlighted that despite the quantitative differences, the three models are consistent in terms of dependence on heterogeneity and of main parameters (e.g. g_CaL_, i_NaKmax_, g_Kr_) importance. Considering that they were developed for different species (human vs. rabbit), from different data and based on different hypotheses (M-vs. Ca^2+^-clock), this is not an obvious result.

### Well-coupled SAN tissues synchronize their behavior despite intercellular heterogeneity

Heterogeneity is known to be an important feature of the sinus node and is thought to be fundamental to the overall normal pacemaking in the mammalian heart [8,17]. Based on this evidence, in our simulations of tissue heterogeneity, we expect to find that cells can coordinate and synchronize their unique discharge frequency to a common one, the one that dictates the human heartbeat. We confirmed previous findings that showed that in rabbit multicellular simulations cells synchronized their frequency (not shown) [5]. In Figure **6**.6 we demonstrated that this is also true in the human tissue at almost any level of heterogeneity and coupling that we tested. The only exception occurred when the cells were weakly coupled (R equal to 10,000 MΩ) and characterized by a high degree of heterogeneous ionic channel expression (σ equal to 0.4, 0.5). In these extreme scenarios, the low coupling was not sufficient to restore beating to the cells that had become silent due to the heterogeneity. Our simulations show that the cells synchronize their discharge frequency, APA and APD, when strongly coupled (Figure **6**.6B-D). The tissue retained a limited residual variability in APA and APD at σ equal to 0.5.

**Figure 6.**
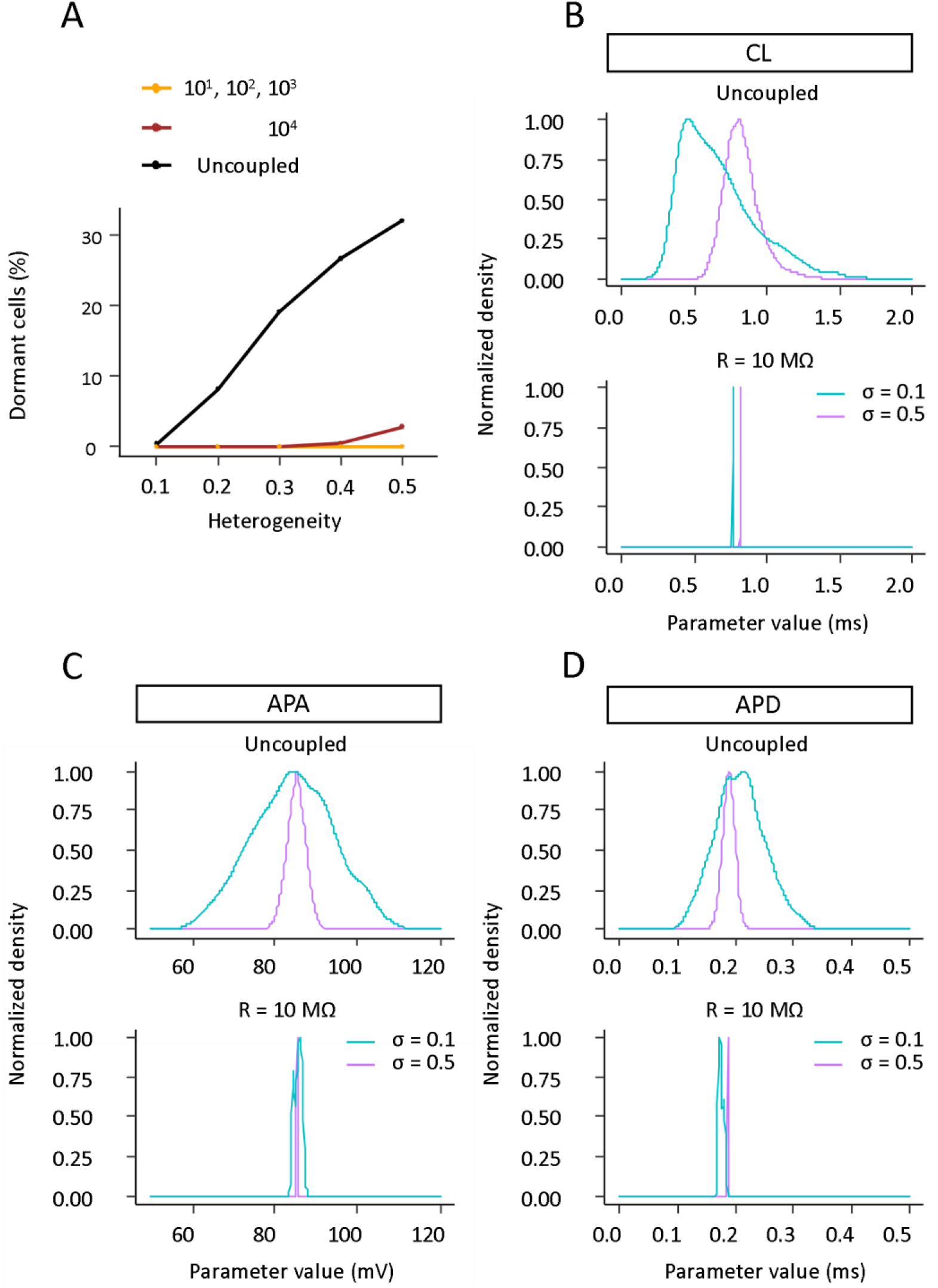
SAN cells synchronize their electrical properties when coupled in a tissue. (A) When coupled together heterogeneous SAN cells give rise to a spontaneously beating tissue. The only exception occurs at very high values of heterogeneity (σ equal to 0.4 and 0.5) and very high levels of intercellular resistance (R equal to 10,000 MΩ). (B-D) The population of cells synchronizes its AP metrics: cycle length (CL), action potential amplitude (APA) and action potential duration (APD) when well connected in a tissue.

### Ionic current perturbations alter the relationship between gap junctional coupling and SAN automaticity

From the previous simulations, we concluded that stronger intercellular coupling favors the synchronization of a monolayer of SAN cells. In the face of physiological heterogeneity that caused some cells to cease beating when isolated, coupling in a tissue restored spontaneous pacemaker functionality to the cells. In addition to the physiological heterogeneity, we then introduced perturbations in the ionic currents that are fundamental for the automaticity of pacemaker cells. Figure 7A shows the impact of diminished I_CaL_ on the single cell AP of the Fabbri model. Blocking g_CaL_ gradually from 10% to 25% causes the cell to beat with a slower frequency and lower AP amplitude. The beating stops once I_CaL_ is blocked by 50%. Next, we analyzed the consequences of the same perturbation on the tissue. Since the latter is heterogeneous, blockade of Ca^2+^ channels within the tissue meant a shift in the distribution of g_CaL_ as shown in Figure 7B. Compared to the baseline tissue, where cellular heterogeneity (σ equal to 0.1) was considered without any other perturbation of individual currents, the tissue where g_CaL_ was blocked by 50% showed a larger fraction of isolated dormant cells and the ability of the residual spontaneously beating cells to drive them varied depending on the degree of coupling. Interestingly, we found that certain levels of intermediate intercellular resistances were beneficial to the SAN tissue automaticity. For instance, Figure 7C depicts the AP generated by a cell within the tissue. When uncoupled and with 50% block of g_CaL_, the cell loses its ability to spontaneously beat. On the other hand, when coupled at intermediate intercellular resistance values, its automaticity (beating capability) is restored. Coupling at lower levels of intercellular resistance causes the cell to stop beating again. Generalizing to the whole tissue, we show in Figure 7D that although 96.6% of isolated cells are incapable of beating spontaneously, under certain degrees of coupling (from 900 MΩ to 4000 MΩ), the same cells coupled in the tissue recover their automaticity. Resistances below or above this range prevent the beating. Tissues where g_CaL_ was blocked by either 10% or 25% are capable of stimulus generation andsynchronization at any levels of coupling (Figure **8**A). The electrical properties of the tissue at different levels of I_CaL_ inhibition and at intermediate coupling resistance (R=1,000 MΩ) are quantified in Figure 7E. Blockade of I_CaL_ up to 25% causes the monolayer of SAN cells to beat at a lower frequency. However, the opposite effect is seen when I_CaL_ is inhibited by 50%.

**Figure 7.**
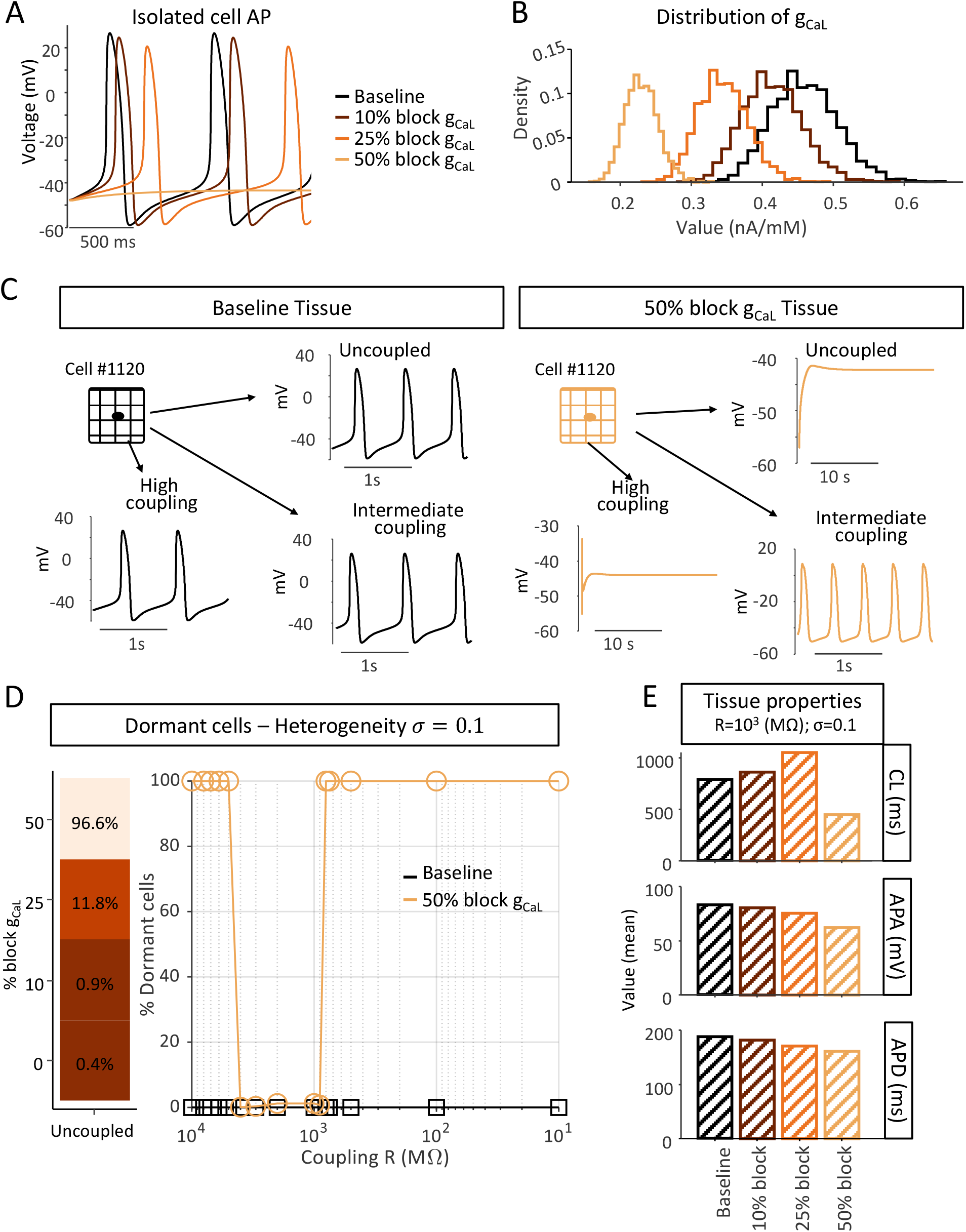
Certain coupling conditions restore automaticity to a prevalently dormant SA node tissue. (A) Effect of Ca^2+^ blockade on Fabbri single cell. (B) Distribution of g_CaL_ in the tissue at varying degrees of Ca^2+^ blockade (cellular heterogeneity factor σ equals 0.1). (C) Comparison of the electrical activity of a cell within the tissue (σ equals 0.1) before and after blockade of Ca^2+^ by 50%. (D) Dormant cells return to beat at intermediate values of coupling. (E) Quantification of the average tissue CL, APA and APD at varying degrees of Ca^2+^ blockade.

**Figure 8.**
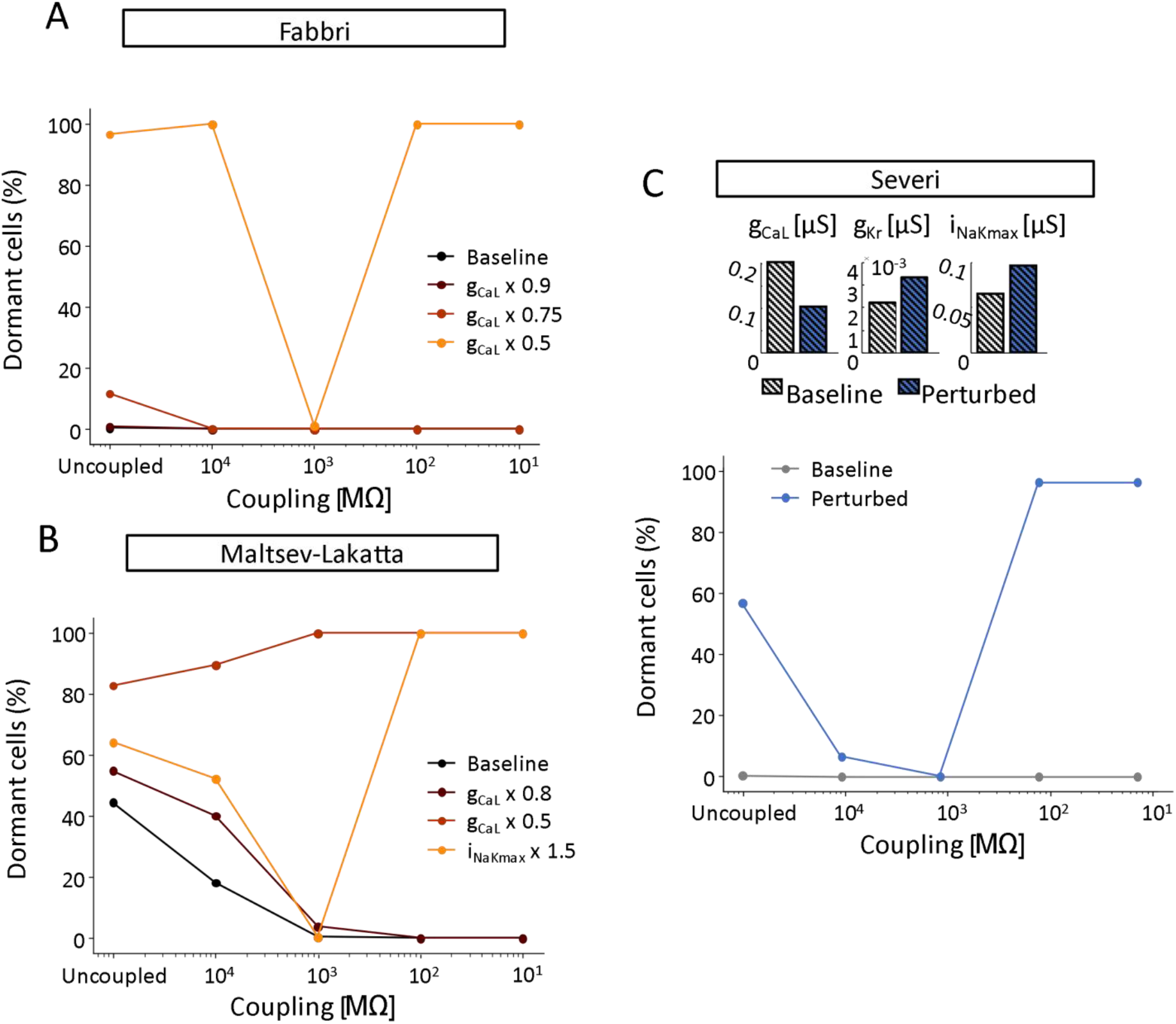
Pathophysiological changes in ionic currents lead to a pattern of tissue automaticity dependent on the degree of intercellular coupling. (A) Effect of L-type Ca^2+^ current (I_CaL_) perturbation in the Fabbri tissue model (σ equal to 0.1). (B) Effect of perturbation in I_CaL_ and Na^+^/K^+^ pump (I_NaK_) in the Maltsev-Lakatta tissue model (σ equal to 0.4). (C) Effect of combined I_CaL_, rapid delayed rectifier K^+^ current (I_Kr_), and I_NaK_ perturbation in the Severi tissue model (σ equal to 0.2).

Moreover, the results presented in Figure **8**B and C suggest that this coupling effect is not limited to perturbations in I_CaL_ nor is it model or species-specific. In fact, certain perturbations involving I_CaL_ and the Na^+^/K^+^ pump (I_NaK_) in the Maltsev-Lakatta model led to a similar trend of coupling effects on the automaticity. For instance, the Severi tissue model showed the same coupling effect when a combination of three currents, rapid delayed rectifier K^+^ current (I_Kr_), I_CaL_ and I_NaK_ were varied. Altogether this suggests that in particular circumstances intermediate coupling can exert a protective effect in response to insults that threaten the normal function of the SAN. Although this phenomenon is seen across models, the specific values of intercellular resistance at which it manifests may vary among models and with the degree of heterogeneity.

### Ca^2+^ channel blockade affects the synchronization of the human sinus node AP phase

To evaluate whether there are further differences between the electrical activity of normal and perturbed tissue we computed a metric similar to the conduction velocity. Given that the concept of conduction velocity is difficult to define in the case of the sinoatrial node, in which cells can synchronize the phase of their action potential in addition to their frequency, we called this metric a delay in activation. We performed this analysis in the case of heterogeneous human tissue (σ = 0.1) and varying magnitude of g_CaL_. In tissues where Ca^2+^ channels were blocked by either 10 or 25% the distribution of activation times was wider compared to the baseline, hence the delay in activation was inversely proportional to the level of I_CaL_. This was compatible with a longer CL and slower beating rate (Figure 7). The activity of the cells was highly asynchronous in the case of I_CaL_ blocked by 50% and the shorter cycle lengths made this same analysis challenging to perform. Activation maps were generated in this case which showed that multiple areas within the tissue seemingly initiated the stimulus at the same time and led to multiple propagating waves (not shown).

### Clusters of beating cells drive AP propagation throughout tissue with reduced excitability

In Figure 7 we saw that when I_CaL_ is inhibited by 50%, less than 4% of cells are still capable of pacemaker activity. We then had placed all of the cells randomly in a matrix representative of a patch of SAN tissue. Here instead we clustered the spontaneously beating cells and placed them in the tissue surrounded by the dormant cells that stopped beating in response to I_CaL_ blockade (9). We compared the effect of coupling on SAN automaticity in these two different tissue configurations. The results (Figure **9**9) suggest that when pacemaker cells form a small cluster, they can drive propagation even in conditions of intercellular coupling that previously prevented excitability. Theoretically, these results might signify that pacemaker cells, co-localized in a subregion of the tissue, were protected by the hyperpolarizing effect of dormant cells. Similar considerations have been proposed to explain the ability of the SAN center to overcome the hyperpolarizing influence of the atrium [18].

**Figure 9.**
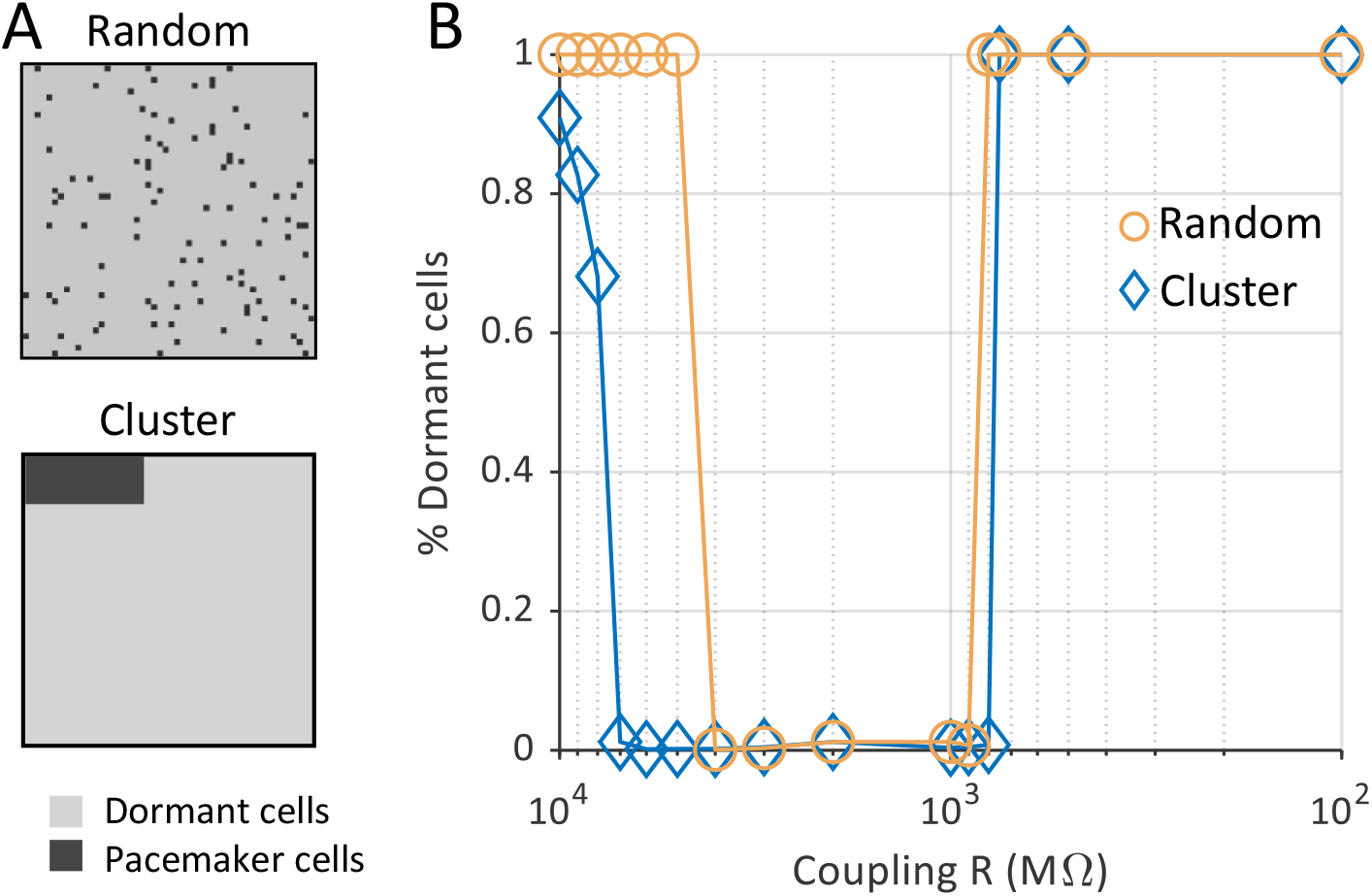
A small cluster of pacemaker cells can drive a prevalently dormant tissue. (A-top) (A) In the random tissue configuration dormant and pacemaker cells are interspersed in the matrix. (A-bottom) In the cluster configuration pacemaker cells are confined to a small portion of the matrix surrounded by dormant cells. Here dormant cells are cells that fail to depolarize after inhibition of I_CaL_ by 50%. (B) The range of intercellular coupling compatible with AP generation and entrainmententrainment, i.e. reduced percentage of dormant cells, is wider in the cluster tissue configuration compared to the random. Results shown here were obtained with Fabbri human model (σ equal to 0.1).

### Intermediate coupling encourages tissue beating due to interactions between driving cells and dormant cells

Results presented thus far suggest that to understand the mechanisms of excitability in the overall tissue, we need to take a closer look at what occurs in the vicinity of the few pacemaker cells present in the tissue. In particular, we are interested in uncovering how, under conditions when a majority of cells do not exhibit spontaneous beating, a small percentage of cells is able to drive tissue depolarization within a narrow range of intercellular coupling.

To investigate this question, we performed simulations with a spontaneously beating and a dormant cell (both extracted from the σ = 0.3, g_CaL_ block = 50% tissue). From the simulation results of the two cells, we computed the average I_net_ during the central portion of the DD and, when action potentials occurred, I_net_ at the TOP (Figure **50**A). Plots of these quantities over a range of coupling resistances help to explain why the spontaneously beating cell (Cell 1) is only able to drive the dormant cell (Cell 2) at intermediate coupling values (Figure **60**B). When coupling between the two cells is strong (R = 10^1^-10^2^ MΩ), the dormant cell can suppress action potentials in the cell that would otherwise beat spontaneously (Figure **70**C, right). This occurs because the large gap junctional current through the low resistance junction results in a small magnitude of diastolic I_net_ in both cells. Under these conditions, TOP I_net_ is undefined since neither cell reaches TOP. With reduced coupling between the two cells (R = 10^3^ MΩ) gap junctional current between the two cells is reduced, which allows a larger magnitude of diastolic I_net_ in Cell 1. This enables Cell 1 to reach its TOP and fully activate its inward current, thereforeit is able to drive the otherwise dormant Cell 2 (Figure **80**C, middle). Finally, when coupling between the cells is reduced further (R = 10^4^ and higher), a large inward diastolic I_net_ in Cell 1 is able to bring this cell to TOP, but the small magnitude of coupling current means that Cell 1 is unable to drive beating in Cell. Thus, intermediate values of coupling represent a “sweet spot” at which the spontaneously beating cell and the dormant cell can be synchronized.

To further support this view, the same analysis of I_net_ was applied to the whole 2D tissue (σ = 0.3, g_CaL_ block = 50%). Being this a significantly more complex situation, it was necessary to divide the cells into categories based on their behavior, as explained in the Methods section. As an additional simplification, in these simulations the last MDP of spontaneous cells in the uncoupled condition was set as their initial condition, in order to obtain a better-defined diastolic phase. For dormant cells, the most depolarized MDP of the spontaneous cells was used as the initial condition.

As with the cell pair, when coupling between cells is strong (R = 10^1^-10^2^ MΩ), dormant cells suppress electrical activity in spontaneous cells (Figure **10** C, right) by draining current during the diastolic phase. Thus, TOP is not reached and the entirety of the tissue becomes “stopped” or “dormant”. With reduced coupling (R = 10^3^ MΩ), spontaneous cells retain a larger fraction of diastolic I_net_ which allows them to reach the TOP. The first cells doing so are called “driving”, since they generate an inward current which is large enough (Figure **10** B, lower panel) to drive the dormant ones (Figure **10** C, middle). Further reducing the coupling allows more cells to reach the TOP on their own (Figure **10** A), but on the other hand it hampers their ability to drive the rest of the tissue because of a reduced gap junctional current (R = 10^4^ and higher).

**Figure 10.**
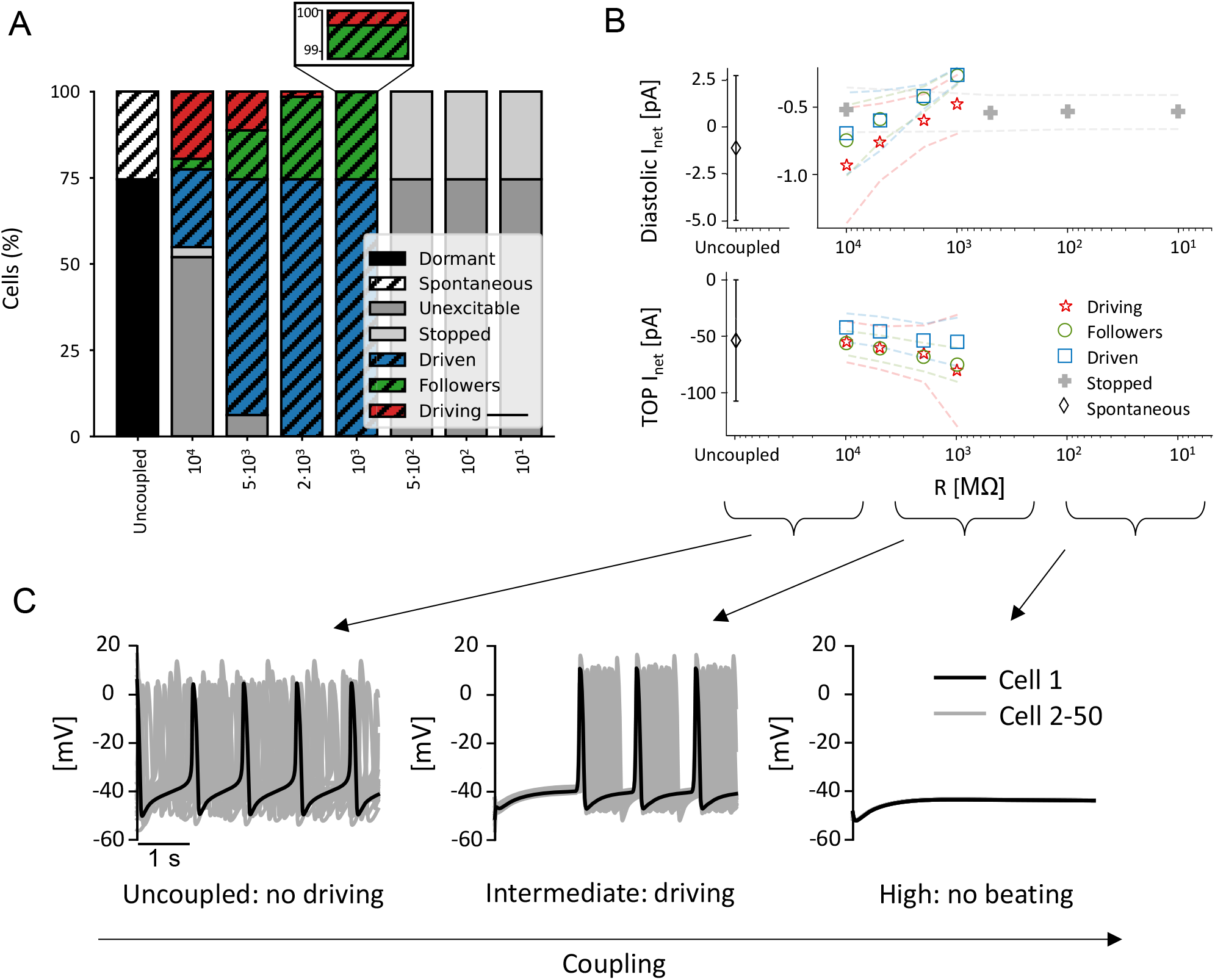
Coupling spontaneous cells with dormant cells inside a tissue. (A) Percentages of cells composing each category at different degrees of cellular coupling. (B) I_net_ trends during diastole (top) and at TOP (bottom) with respect to different degrees of cellular coupling for every cell category. Average value ± standard deviation. (C) Electrical activity of 50 cells (4^th^ column of the 2D tissue matrix) when they are coupled with different intercellular resistances.

**Figure 90.**
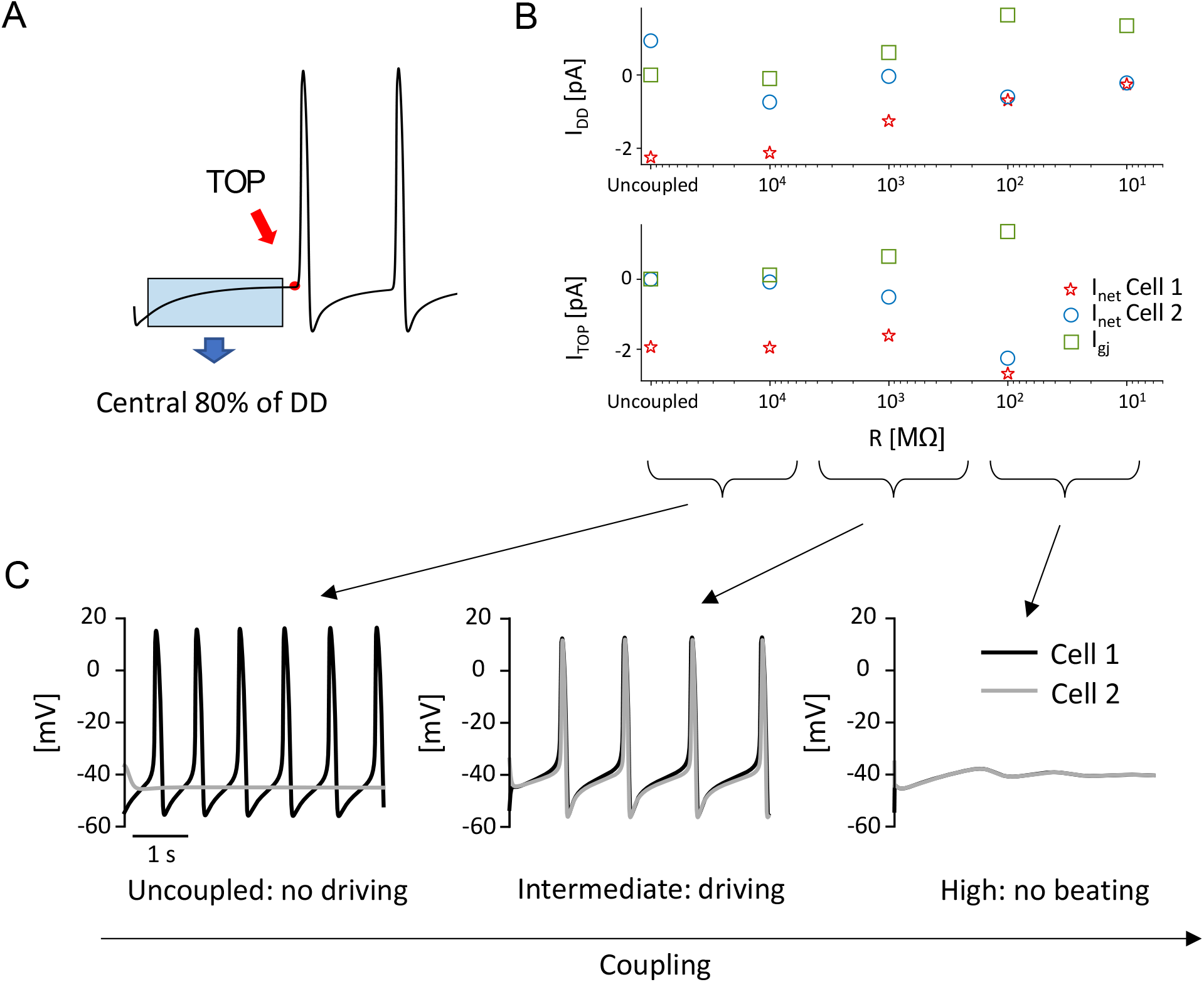
Coupling a spontaneous cell with a dormant cell. (A) Average I_net_ and I_gj_ were extracted from the central 80% portion of the first occurrence of DD (from the beginning of the simulation to the first TOP); TOP I_net_ and I_gj_ were extracted at TOP instant. (B) I_net_ and I_gj_ trends during diastole (top) and at TOP (bottom) with respect to different degrees of cellular coupling for the two cells. I_gj_ is plotted in green for Cell1 (for Cell 2 it is the same with the opposite sign). (C) Behavior of the two cells depending on coupling: both cells are beating only for intermediate coupling values.

## DISCUSSION

In the present study, we investigated the effect of different levels of cellular heterogeneity and intercellular coupling on human and rabbit SAN pacemaking. We simulated both healthy and diseased conditions, highlighting the remarkable robustness of this tissue. Results showed that although increased cellular heterogeneity leads to a growing fraction of cells losing automaticity, coupling between cells allows for rhythmic activity in the whole tissue. Of note, this remained true for nearly all combinataions of heterogeneity and coupling. When we simulated diseased conditions by increasing or decreasing levels of fundamental ionic currents, rhythmic depolarizations of the tissue could fail. However, even under these extreme conditions, intermediate values of gap-junctional resistance could rescue SAN electrical activity, and simulations provided mechanistic insight into this unusual phenomenon. This behavior was seen in all 3 models that we examined, [9][10][11], suggesting that it may be a general property of SA nodal entrainment, rather than specific to an individual model.

### Comparison with previous computational SAN studies

Mathematical modeling has been employed as a tool to understand the mechanisms of sinoatrial pacemaker coupling and entrainment for more than two decades. Early studies, such as by Michaels et al. (1987) and [19], demonstrated how simulations of SAN pacemaker activity in models of coupled cells can provide insights and encourage new hypotheses about cardiac electrical conduction. Mathematical modeling, in conjunction with animal experiments, has been instrumental in understanding that the heartbeat is likely to be dictated by the mutual entrainment of multiple spontaneously beating cells that synchronize their activity. Over the years, many investigators developed models to further describe the role of mutual entrainment of heterogeneous cells in the generation of the pacemaker activity. For instance, Oren and Clancy (2010) showed that connection between the SAN and the atrium might be sufficient to impart the different features of peripheral SAN compared with central SAN APs. Conversely, Inada et al. (2014), argued for the necessity of gradual changes in cell size, ionic current densities and intercellular coupling from center to periphery were necessary. In particular, they suggested that the expression of Na_v_1.5 and Cx43 in the periphery of the SAN might be fundamental for driving the propagation to the atrium. Additional relevant insights were obtained by Gratz et al. (2018), who noted complex interactions between ion channel conductances, intercellular coupling, and the synchronization of small SAN tissues. Our work builds upon these studies by assessing synchronization in the human SAN, in both physiological and pathological conditions.

### Modeling insights into the physiology and pathophysiology of the SAN

We simulated the effects on SAN automaticity of both physiological heterogeneity in ionic current densities and pathological changes in ionic currents. The results showed that this heterogeneity is compatible with synchronization within a large monolayer of either human or rabbit SAN cells. Moreover, we suggest that under conditions of reduced coupling between nodal cells, this heterogeneity helps to impart remarkable resilience that allows for stimulus propagation even in the presence of pathological changes in the cellular electrical properties.

Many pathological alterations, both congenital and acquired, can lead to SAN dysfunction. Mutations in ionic channels [21,22] have been discovered that underlie congenital disease, but the acquired forms are far more common and are more prevalent in older individuals. With aging the SAN function changes, frequently presenting as a decrease in heart rate; but also as a decrease in peripheral Na^+^ channel expression [23,24] and in Cx43 expression [25]. Both these changes contribute to slower conduction of the electrical stimulus. Heart failure (HF), chronic atrial fibrillation and cell apoptosis [26] also contribute to structural and electrical remodeling of the sinus node, possibly determining SAN dysfunction.

Whatever the cause leading to a loss of spontaneous beating in isolated SAN cells, our results show that SAN tissue can experience a higher level of robustness. A significant cellular heterogeneity partially compensates for these consequences, guaranteeing that more cells show APs even with a degraded coupling. Our simulations revealed that specific coupling strengths, falling in the range 900-4000 MΩ (Figure 7D), or 0.25-1.1 nS when expressed as conductances, allow the tissue to beat despite blockage of I_CaL_. Comparing this range of coupling strengths to the existing literature, we find that it sits at the low end of previously reported values. For instance, experimental studies on rabbit SAN suggested that 0.5 nS would allow for frequency entrainment and 10 nS for waveform entrainment [3]). Other computational works have employed inter-cellular resistances of 7.5 nS [20] and 25 nS [16], whereas experiments have estimated values such as 0.6-25 nS [15] and 2.6±10 nS [27]. Thus, the protective range of coupling that our simulations identified, which became relevant under simulated pathological conditions, is consistent with the reduced coupling between SAN cells that has been identified under some pathological conditions. One could even speculate that remodeling of gap junctions (a decrease of Cx-expression) under such conditions may halp to protects the SAN from failure.

Naively, one might expect that stronger coupling between SA nodal myocytes will be beneficial, since this will lead to faster propagation and enhanced synchronization of the cells within the node. Although our results are consistent with this idea under normal conditions, our findings also highlight a potential advantage of reduced coupling – namely that this can impart the tissue with greater resilience under conditions that impair spontaneous beating in individual myocytes. Indeed, it is remarkable that under particular conditions, fewer than 10% of the cells in the tissue can drive electrical activity in the remaining 90% of myocytes that do not.

### A possible explanation for the protective role of intermediate coupling: AP vs. DD intercellular interactions

To attempt to explain the protective range of coupling strengths under pathological conditions, (Figures 7-8), we formulated a hypothesis based on the concepts of tonic and phasic entrainment well established in SAN literature (Michaels et al. 1986; Verheijk et al., 1998). The new aspect here is that in our simulations these two types of interaction not only regulate SAN synchronism, but also determine the presence of spontaneous beating inside the tissue. In other words, spontaneous cells manage to drive dormant cells only if two conditions are satisfied. First, spontaneous cells have to reach the take-off potential. Second, they have to supply enough current to the neighboring dormant cells. Coupling resistance is, in this condition, the most critical parameter: if too low, dormant cells will hyperpolarize the spontaneous ones during the DD phase, preventing them from reaching the threshold for AP firing. On the other hand, if coupling resistance is too high, spontaneous cells will not manage to supply enough current during their AP to depolarize dormant cells. In either case, no electrical activity can be seen in the tissue. However, intermediate values of coupling guarantee that both conditions are satisfied. During diastole, when the voltage difference is low, I_gj_ is negligible, whereas during the upstroke, I_gj_ increases and allows dormant cells to depolarize. Figure **7** elucidates this mechanism for a simple cell pair, which remains valid in tissue simulations, as shown in Figure **8**.

### Are dormant cells present inside the sinoatrial node?

Recently, a combined experimental and computational work [8] reported that about half of SAN cells isolated from guinea pig hearts did not show APs, but did so in response to isoprotenerol. This behaviour was explained in terms of entrainment between membrane and calcium clocks. Considering that I_CaL_ has an important impact on clock entrainment, our human model is consistent with this work in showing that a progressive reduction in this current leads to an increasing percentage of isolated dormant cells. However, we further demonstrate that when these silent cells are coupled with a minority of spontaneous cells, the tissue can exhibit stable electrical activity without the need of a basal sympathetic stimulation. Our results also suggest that substantial amounts of isolated dormant cells should not be regarded as a consequence of the isolation procedure but could be actually present inside the SAN, being compatible with its physiological behaviour. The excessive presence of dormant cells poses nevertheless a threat to SAN function, restricting the coupling range in which rhythmic electrical activity can be appreciated. This highlights the perils of pathologies that depress SAN single cell activity, like SAN dysfunction.

### Limitations and future developments

In this work, we used a modeling approach that allowed us to study a monolayer of SAN cells where we varied the degree of cellular heterogeneity and intercellular coupling. This strategy offered us the possibility of investigating tissue automaticity and synchronization mechanisms that would be impossible to model at the single-cell level. Several limitations of our approach, however, should be mentioned. First, the cellular heterogeneity was represented as random differences in ion channel expression between cells, and we did not consider gradients across the tissue in cell type, size, or shape. Additionally, we did not take into account the presence of non-pacemaker cells interspersed in the SAN such as fibroblasts, atrial cells and adipocytes [28]. Another structure simplification is the idealized geometry represented by a square sheet, far from the 3D banana-shaped anatomy of the SAN [29,30]. Our tissue, which comprised 2500 cells, is comparable to the number of cells (about 5000) from which the stimulus is believed to originate from in rabbit SAN [31], but represents only a fraction of the human SAN. These limitations can all be addressed in future simulation studies that can shed additional light on mechanisms of SAN pacemaking.

## Conclusions

In conclusion, we have shown how multiscale mathematical modeling can be used to gain insight into the importance of cellular heterogeneity and intercellular coupling for efficacious cardiac entrainment. Previous multi-cellular studies have shown that synchronization of heterogenous cells is responsible for the sinoatrial node pacemaker function in rabbits. Our data confirmed that the same phenomenon occurs in a two-dimensional model of the human sinoatrial node. In addition, our study suggests that certain degrees of intercellular coupling make the sinoatrial node resistant to ionic perturbations that might be provoked by mutations and/or drug therapies.

## Supplementary Material

**Table 1.** Hardware and software specification for model reproducibility.

| **Hardware** |  |  |  |
| --- | --- | --- | --- |
| **Workstation 1** |  | *Workstation 2* |  |
| **Operating system** | Ubuntu 19.04 | *Operating system* | Windows 10 |
| **RAM** | 64.0 GB | *RAM* | 16.0 GB |
| **CPU** | 16-core AMD Ryzen threadripper 2950x | *CPU* | Intel® Core™ i7-8700K |
| **GPU** | 12 GB Nvidia Titan V | *GPU* | NVIDIA GeForce GTX 1060 6 GB |
| **Software** |  |  |  |
| **Workstation 1** |  | *Workstation 2* |  |
| **Simulation** | MATLAB R2019b;<br>MATLAB GPU coder | *Simulation* | MATLAB R2020a;<br>CUDA 8.0; Visual Studio 2015 |
| **Integration** | Euler method (fixed step of 10 μs) | *Integration* | Euler method (fixed step of 10 μs) |
| **Analysis** | MATLAB R2019b<br>Python 3.7 | *Analysis* | MATLAB R2020a<br>R-4.0.3 |

## Bibliography

1. Brennan JA, Chen Q, Gams A, Dyavanapalli J, Mendelowitz D, Peng W, et al. Evidence of Superior and Inferior Sinoatrial Nodes in the Mammalian Heart. JACC Clin Electrophysiol. 2020;6: 1827–1840. doi:10.1016/j.jacep.2020.09.012

2. Jalife J. Mutual entrainment and electrical coupling as mechanisms for synchronous firing of rabbit sino-atrial pace-maker cells. J Physiol. 1984;356: 221–243. doi:https://doi.org/10.1113/jphysiol.1984.sp015461

3. Verheijck EE, Wilders R, Joyner RW, Golod DA, Kumar R, Jongsma HJ, et al. Pacemaker synchronization of electrically coupled rabbit sinoatrial node cells. J Gen Physiol. 1998;111: 95–112. doi:10.1085/jgp.111.1.95

4. Ypey DL, Clapham DE, DeHaan RL. Development of electrical coupling and action potential synchrony between paired aggregates of embryonic heart cells. J Membr Biol. 1979;51: 75– 96. doi:10.1007/BF01869344

5. Gratz D, Onal B, Dalic A, Hund TJ. Synchronization of Pacemaking in the Sinoatrial Node: A Mathematical Modeling Study. Front Phys. 2018;6. doi:10.3389/fphy.2018.00063

6. Mata AN, Alonso GR, Garza GL, Fernández JRG, García MAC, Ábrego NPC. Parallel simulation of the synchronization of heterogeneous cells in the sinoatrial node. Concurr Comput Pract Exp. 2020;32: e5317. doi:https://doi.org/10.1002/cpe.5317

7. Michaels DC, Matyas EP, Jalife J. Mechanisms of sinoatrial pacemaker synchronization: A new hypothesis. Circ Res. 1987;61: 704–714. doi:10.1161/01.RES.61.5.704

8. Kim MS, Maltsev AV, Monfredi O, Maltseva LA, Wirth A, Florio MC, et al. Heterogeneity of calcium clock functions in dormant, dysrhythmically and rhythmically firing single pacemaker cells isolated from SA node. Cell Calcium. 2018;74: 168–179. doi:10.1016/j.ceca.2018.07.002

9. Maltsev VA, Lakatta EG. Synergism of coupled subsarcolemmal Ca2+ clocks and sarcolemmal voltage clocks confers robust and flexible pacemaker function in a novel pacemaker cell model. Am J Physiol Heart Circ Physiol. 2009;296: H594–615. doi:10.1152/ajpheart.01118.2008

10. Severi S, Fantini M, Charawi LA, DiFrancesco D. An updated computational model of rabbit sinoatrial action potential to investigate the mechanisms of heart rate modulation. J Physiol. 2012;590: 4483–4499. doi:10.1113/jphysiol.2012.229435

11. Fabbri A, Fantini M, Wilders R, Severi S. Computational analysis of the human sinus node action potential: model development and effects of mutations. J Physiol. 2017;595: 2365– 2396. doi:10.1113/JP273259

12. Sarkar AX, Sobie EA. Quantification of repolarization reserve to understand interpatient variability in the response to proarrhythmic drugs: a computational analysis. Heart Rhythm. 2011;8: 1749–1755. doi:10.1016/j.hrthm.2011.05.023

13. Sobie EA. Parameter Sensitivity Analysis in Electrophysiological Models Using Multivariable Regression. Biophys J. 2009;96: 1264–1274. doi:10.1016/j.bpj.2008.10.056

14. Lee Y-S, Liu OZ, Hwang HS, Knollmann BC, Sobie EA. Parameter sensitivity analysis of stochastic models provides insights into cardiac calcium sparks. Biophys J. 2013;104: 1142– 1150. doi:10.1016/j.bpj.2012.12.055

15. Verheule S, van Kempen MJ, Postma S, Rook MB, Jongsma HJ. Gap junctions in the rabbit sinoatrial node. Am J Physiol Heart Circ Physiol. 2001;280: H2103–2115. doi:10.1152/ajpheart.2001.280.5.H2103

16. Inada S, Zhang H, Tellez JO, Shibata N, Nakazawa K, Kamiya K, et al. Importance of Gradients in Membrane Properties and Electrical Coupling in Sinoatrial Node Pacing. PLoS ONE. 2014;9. doi:10.1371/journal.pone.0094565

17. Monfredi O, Dobrzynski H, Mondal T, Boyett MR, Morris GM. The Anatomy and Physiology of the Sinoatrial Node—A Contemporary Review. Pacing Clin Electrophysiol. 2010;33: 1392– 1406. doi:https://doi.org/10.1111/j.1540-8159.2010.02838.x

18. Dobrzynski H, Li J, Tellez J, Greener ID, Nikolski VP, Wright SE, et al. Computer three-dimensional reconstruction of the sinoatrial node. Circulation. 2005;111: 846–854. doi:10.1161/01.CIR.0000152100.04087.DB

19. Joyner RW, van Capelle FJ. Propagation through electrically coupled cells. How a small SA node drives a large atrium. Biophys J. 1986;50: 1157–1164. doi:10.1016/S0006-3495(86)83559-7

20. Oren RV, Clancy CE. Determinants of heterogeneity, excitation and conduction in the sinoatrial node: a model study. PLoS Comput Biol. 2010;6: e1001041. doi:10.1371/journal.pcbi.1001041

21. Butters TD, Aslanidi OV, Inada S, Boyett MR, Hancox JC, Lei M, et al. Mechanistic links between Na+ channel (SCN5A) mutations and impaired cardiac pacemaking in sick sinus syndrome. Circ Res. 2010;107: 126–137. doi:10.1161/CIRCRESAHA.110.219949

22. Choudhury M, Boyett MR, Morris GM. Biology of the Sinus Node and its Disease. Arrhythmia Electrophysiol Rev. 2015;4: 28–34. doi:10.15420/aer.2015.4.1.28

23. Alings AM, Bouman LN. Electrophysiology of the ageing rabbit and cat sinoatrial node--a comparative study. Eur Heart J. 1993;14: 1278–1288. doi:10.1093/eurheartj/14.9.1278

24. Tellez JO, Mączewski M, Yanni J, Sutyagin P, Mackiewicz U, Atkinson A, et al. Ageing-dependent remodelling of ion channel and Ca2+ clock genes underlying sino-atrial node pacemaking. Exp Physiol. 2011;96: 1163–1178. doi:https://doi.org/10.1113/expphysiol.2011.057752

25. Jones SA, Lancaster MK, Boyett MR. Ageing-related changes of connexins and conduction within the sinoatrial node. J Physiol. 2004;560: 429–437. doi:https://doi.org/10.1113/jphysiol.2004.072108

26. Swaminathan PD, Purohit A, Soni S, Voigt N, Singh MV, Glukhov AV, et al. Oxidized CaMKII causes cardiac sinus node dysfunction in mice. J Clin Invest. 2011;121: 3277–3288. doi:10.1172/JCI57833

27. Anumonwo J M, Wang H Z, Trabka-Janik E, Dunham B, Veenstra R D, Delmar M, et al. Gap junctional channels in adult mammalian sinus nodal cells. Immunolocalization and electrophysiology. Circ Res. 1992;71: 229–239. doi:10.1161/01.RES.71.2.229

28. John RM, Kumar S. Sinus Node and Atrial Arrhythmias. Circulation. 2016;133: 1892–1900. doi:10.1161/CIRCULATIONAHA.116.018011

29. Csepe TA, Zhao J, Hansen BJ, Li N, Sul LV, Lim P, et al. Human Sinoatrial Node Structure: 3D Microanatomy of Sinoatrial Conduction Pathways. Prog Biophys Mol Biol. 2016;120: 164–178. doi:10.1016/j.pbiomolbio.2015.12.011

30. Dobrzynski H, Anderson RH, Atkinson A, Borbas Z, D’Souza A, Fraser JF, et al. Structure, function and clinical relevance of the cardiac conduction system, including the atrioventricular ring and outflow tract tissues. Pharmacol Ther. 2013;139: 260–288. doi:10.1016/j.pharmthera.2013.04.010

31. Bleeker WK, Mackaay AJ, Masson-Pévet M, Bouman LN, Becker AE. Functional and morphological organization of the rabbit sinus node. Circ Res. 1980;46: 11–22. doi:10.1161/01.RES.46.1.11

